# A comparative study of isothermal nucleic acid amplification methods for SARS-CoV-2 detection at point-of-care

**DOI:** 10.1101/2020.05.24.113423

**Authors:** Diem Hong Tran, Hoang Quoc Cuong, Hau Thi Tran, Uyen Phuong Le, Hoang Dang Khoa Do, Le Minh Bui, Nguyen Duc Hai, Hoang Thuy Linh, Nguyen Thi Thanh Thao, Nguyen Hoang Anh, Nguyen Trung Hieu, Cao Minh Thang, Van Van Vu, Huong Thi Thu Phung

**Affiliations:** NTT Hi-Tech Institute, Nguyen Tat Thanh University, Ho Chi Minh City, Vietnam; Directorial Board, Pasteur Institute in Ho Chi Minh City, Vietnam; Planning Division, Pasteur Institute in Ho Chi Minh City, Vietnam; Medical Analysis Department, Pasteur Institute in Ho Chi Minh City, Vietnam; Microbiology and Immunology Department, Pasteur Institute in Ho Chi Minh City, Vietnam

**Keywords:** SARS-CoV-2, nucleic acid amplification test, LAMP, CPA, PSR, colorimetric, lyophilized kit

## Abstract

COVID-19, caused by the novel coronavirus SARS-CoV-2, has spread worldwide and put most of the world under lockdown. Despite that there have been emergently approved vaccines for SARS-CoV-2, COVID-19 cases, hospitalizations, and deaths have remained rising. Thus, rapid diagnosis and necessary public health measures are still key parts to contain the pandemic. In this study, the colorimetric isothermal nucleic acid amplification tests (iNAATs) for SARS-CoV-2 detection based on loop-mediated isothermal amplification (LAMP), cross-priming amplification (CPA), and polymerase spiral reaction (PSR) were designed and evaluated. The three methods showed the same limit of detection (LOD) value of 1 copy of the targeted gene per reaction. However, for the direct detection of SARS-CoV-2 genomic-RNA, LAMP outperformed both CPA and PSR, exhibiting the LOD value of roughly 43.14 genome copies/reaction. The results can be read with the naked eye within 45 minutes, without cross-reactivity to closely related coronaviruses. Moreover, the direct detection of SARS-CoV-2 RNA in simulated patient specimens by iNAATs was also successful. Finally, the ready-to-use lyophilized reagents for LAMP reactions were shown to maintain the sensitivity and LOD value of the liquid assays. The results indicate that the colorimetric lyophilized LAMP kit developed herein is highly suitable for detecting SARS-CoV-2 nucleic acids at point-of-care.

## INTRODUCTION

Coronavirus is a large family of RNA viruses, including the human coronaviruses which often cause respiratory illnesses with mild cold symptoms [1, 2]. The two exceptions that cause severe diseases include the fatal Severe Acute Respiratory Syndrome Coronavirus (SARS-CoV) [3] and the Middle East Respiratory Syndrome Coronavirus (MERS-CoV) [4]. By February 2020, the mortal pneumonia disease caused by a novel coronavirus called SARS-CoV-2 was named COVID-19 by the World Health Organization (WHO) [5]. In March 2020, the WHO classified the COVID-19 outbreak as “Global Pandemic” [6]. As of February 1^st^, 2021, COVID-19 has spread to over 220 countries and regions worldwide with more than 100 million confirmed cases and 2.2 million casualties [7].

The genome of SARS-CoV-2 is a single-stranded positive-sense RNA molecule that is around 29.9 kb in length [8, 9]. The viral genome is composed of 11 Open reading frames (ORFs), namely *ORF1ab, ORF2* (Spike protein or S gene), *ORF3a*, *ORF4* (Envelope protein or E gene), *ORF5* (Membrane protein or M gene), *ORF6, ORF7a, ORF7b, ORF8, ORF9* (Nucleocapsid protein or N gene), and *ORF10*[10]. The *ORF1ab* gene expresses a polyprotein comprising of 16 nonstructural proteins [10]. The sequences of four structural proteins, including spike (S), envelope (E), membrane (M), and nucleocapsid (N) proteins, share high similarity to those of SARS-CoV and MERS-CoV [9].

The diagnostic standard of SARS-CoV-2 involves clinical symptoms and molecular methods. For an accurate diagnosis, the quantitative reverse transcription PCR (qRT-PCR) for the detection of SARS-CoV-2 was soon introduced by WHO [11]. Currently, the qRT-PCR test, which is carried out using respiratory samples (nasopharyngeal or oropharyngeal swabs), has been widely used as the gold standard method for SARS-CoV-2 diagnosis. Nevertheless, qRT-PCR requires high-cost equipment, and results are only available within a few hours to 2 days, limiting its application at resource-limited settings.

Distinctive virological and serological assays for rapidly detecting SARS-CoV-2 at point-of-care (POC) have been introduced. Virological diagnosis directly detects the viral nucleic acids via isothermal nucleic acid amplification tests (iNAATs) [12–17] and CRISPR-Cas12 based method [18], while serological tests detect the rising titers of antibody between acute and convalescent stages of infection or detect IgM in primary infection [19–22]. However, serological diagnosis usually shows lower sensitivity especially in the early stage of infection [23]. Meanwhile, like PCR, iNAATs that amplify the viral nucleic acids at a constant temperature are expected to determine the presence of infectious viruses even in asymptomatic patients. Loop-mediated isothermal amplification (LAMP) was invented in 2000 and broadly utilized nowadays [24–27]. LAMP was shown to be rapid, specific, and remarkably sensitive compared to conventional PCR [28, 29]. Different findings revealed that the performance of LAMP assays was in a good correlation with qRT-PCR results when evaluated with clinical samples [30–33]. Standard LAMP uses only a *Bst* DNA polymerase possessing strand displacement activity and no modified/labelled DNA probes are required, simplifying the preparation procedure and significantly reducing the cost [34]. Afterward, other isothermal DNA amplification methods developed later, which also depend on the strand displacement activity of the *Bst* DNA polymerase, include cross-priming amplification (CPA) [35, 36] and polymerase spiral reaction (PSR) [37]. While LAMP requires two to three primer pairs, PSR needs only one primer pair, and CPA uses six to eight cross-linked primers. Various studies indicated that PSR and CPA performance regarding sensitivity and specificity was comparable to that of LAMP [38–42]. Simple and fast detection methods for the amplified nucleic acids, such as using SyBr Green dye or pH-sensitive indicators, have been well developed and readily available to visualize the outcome of the three assays [43]. Also, with mutual advantages, including easy operation, fast reaction, low-cost requirements, these methods are suitable diagnostic tools for resource-restricted settings.

Currently, only less than 10% of COVID-19 vaccine candidates have been undergone Phase 3 clinical trials [44]. Among them, preliminary results from three candidates developed by Pfizer, Moderna and AstraZeneca have shown efficacies of or higher than 90% against the development of symptomatic COVID-19 [45–47]. Although all vaccine candidates are considered as still in the early testing stages, many governments have granted emergency authorization of the COVID-19 vaccines. Nevertheless, COVID-19 daily cases and deaths worldwide have kept surging even after vaccine programs have been rolled out across many countries. Additionally, there has been a recent emergence of new COVID-19 variants that are able of transmitting more easily. Together with a global shortage of COVID-19 vaccine supply, the outbreak could even worsen until when effective vaccines are widely distributed. Thus, basic preventative measures including physical distancing, wearing masks, handwashing, and rapid diagnosis of COVID-19 patients as many as possible are still the best tools to defeat SARS-CoV-2, especially in the low- and middle-income countries where the opportunities to access COVID-19 vaccines have still been very far away [48].

In this study, we designed and compared colorimetric LAMP, CPA, and PSR assays for SARS-CoV-2 nucleic acid detection utilizing a pH-sensitive dye for readout visualization. In particular, the options of direct detection of SARS-CoV-2 from clinical samples and the use of ready-to-use lyophilized LAMP were also examined and discussed.

## MATERIALS AND METHODS

### Primer design

Primers for LAMP, CPA, and PSR assays targeting the *N* and *ORF1ab* sequences of SARS-CoV-2 (GenBank accession number MN908947) were designed using the free online software Primer Explorer V5 (https://primerexplorer.jp/e/). Primer selection was carried out as instructed (https://primerexplorer.jp/e/v4_manual/). Two sets of primer pairs for LAMP (targeting the *N* and *ORF1ab* genes), two sets of primer pairs for PSR (targeting the *N* and *ORF1ab* genes), and one set of primer pairs for CPA (targeting the *ORF1ab* gene) were selected. Primers were synthesized by Phu Sa Biochem (Can Tho, Vietnam) and their sequences are listed in Table 1.

**Table 1.**
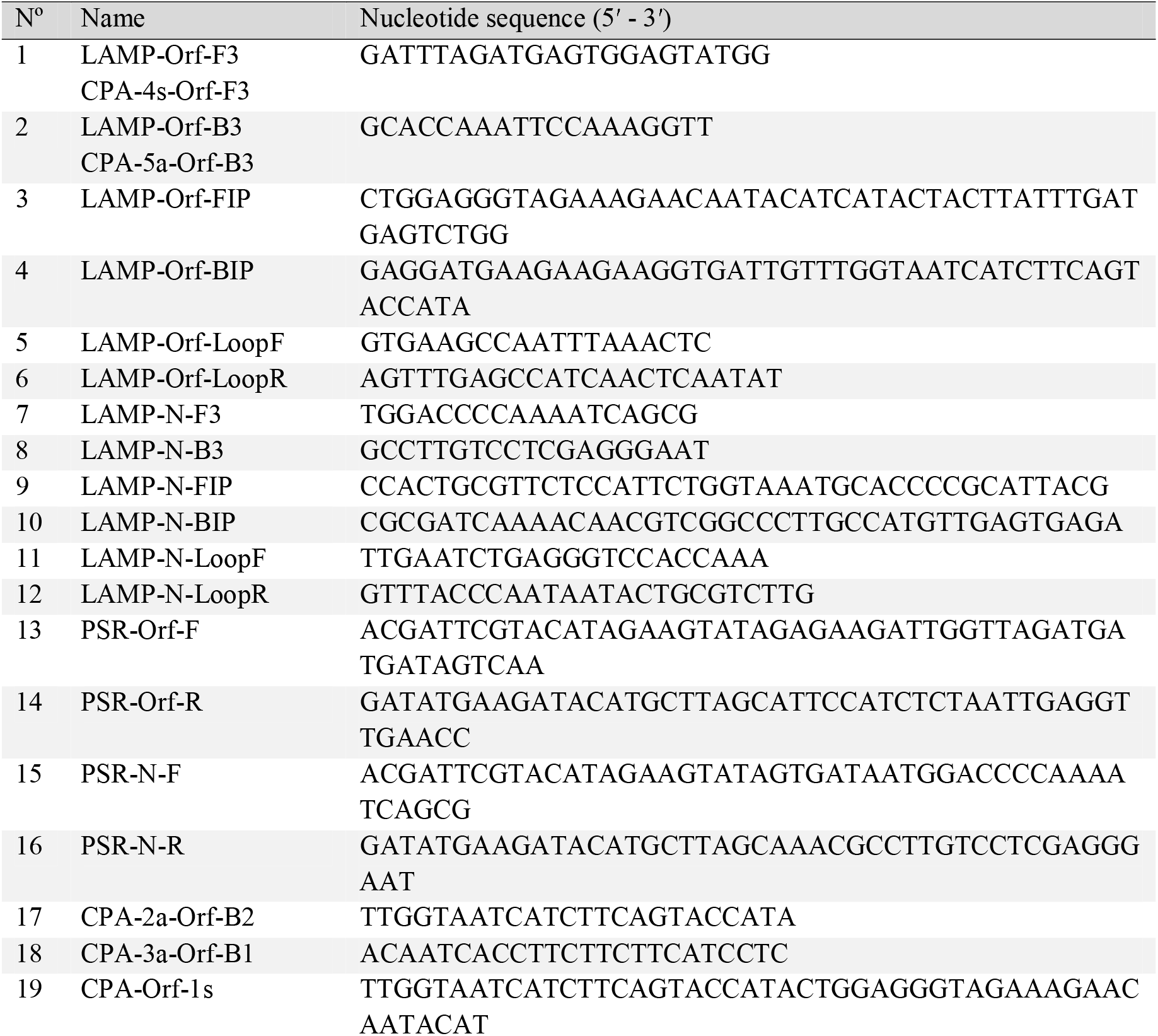
Primers sequences used in this study

### Synthesized DNA template preparation

The synthesized DNA templates were obtained from Phu Sa Biochem, (Can Tho, Vietnam). The sequences from 28274 to 28516 of the *N* gene and from 2853 to 3452 of the *ORF1ab* gene of SARS-CoV-2 (GenBank MN908947) were selected to serve as the positive control templates for iNAATs. The *N* and *ORF1ab* sequences (single-strand) of MERS-CoV (GenBank JX869059.2; 28567-28751 and 3075-3303), SARS-CoV (GenBank MK062184.1, 28278-28482 and 2954-3162), and bat SARS-like-CoV (GenBank MN996532.1, 28205-28504 and MG772933.1, 2851-3350) were also prepared for primer specificity testing.

### Primer specificity analysis

The reference genomes of SARS-CoV-2 and related species were downloaded from NCBI (https://www.ncbi.nlm.nih.gov). The primer sequences were aligned to genomes of different coronaviruses to calculate the number of mismatches using Geneious Prime 2020.0.3 (https://www.geneious.com). The percentage of mismatch was calculated by dividing the total number of different bases between primers and genome sequences by the total length of the primers. The software FastPCR available at http://primerdigital.com/fastpcr.html was used for *in silico* PCR analysis. The software eLAMP downloaded at https://www.nybg.org/files/scientists/dlittle/eLAMP.html was utilized for virtual LAMP analysis.

### Colorimetric iNAATs

WarmStart^®^ Colorimetric LAMP 2X Master Mix (DNA & RNA) was purchased from NEB (MA, USA). The iNAAT reaction volume was 15 μl, consisting of 1 or 5 μl of template sample and 7.5 μl of the WarmStart^®^ Colorimetric Mastermix. The LAMP reaction contains 0.8 μM each inner primer (FIP and BIP), 0.1 μM each outer primer (F3 and B3), 0.2 μM each loop primer (FLoop and Bloop). The CPA reaction contains 0.5 μM cross primer 1s, 0.3 μM each of primers 3a and 2a, 0.05 μM each of displacement primers 4s and 5a. The PSR reaction contains 1.6 μM each primer (PSR-F and PSR-R). The iNAAT reactions were run in BioSan Dry block thermostat Bio TDB-100 for 30 to 45 min at 60 °C (LAMP reaction) or 63 °C (CPA and PSR reactions). The amplification products were visualized by the color shifting from red to yellow of the test reaction, which is based on the use of phenol red, a pH-sensitive indicator as instructed by the manufacturer.

### Evaluation of limit of detection (LOD) of colorimetric iNAATs

The copy number of the synthetic DNA template was calculated using the Endmemo program (http://endmemo.com/bio/dnacopynum.php). The synthesized DNA template was serially diluted to various concentrations and 1 μl of the diluted DNA sample was added to the iNAAT reactions. The reactions were then incubated for 30 min at different temperatures according to each method.

SARS-CoV-2 was isolated from the clinical positive COVID-19 samples and cultured at Pasteur Institute (Ho Chi Minh City, Vietnam). Extracted genomic-RNA of SARS-CoV-2 was quantified via a standard curve based on qRT-PCR Ct-value [49] and then serially diluted. Five μl of the diluted RNA samples were added to the iNAAT reactions. The reactions were then incubated for 45 min at the proper temperature.

### Collection and preparation of clinical specimens

Four nasopharyngeal and 6 oropharyngeal swab specimens from volunteer nurses and doctors were collected at a local hospital (Ho Chi Minh City, Vietnam). All participants were confirmed negative with SARS-CoV-2 by RT-PCR provided their informed consent to participate in the trial. The oropharyngeal or nasopharyngeal specimen was collected using a sterile flocked plastic swab which was then soaked into 400 μl of nuclease-free water. Fresh samples were kept on ice until analysis or frozen for subsequent assays.

To prepare the simulated clinical specimens, various concentrations of SARS-CoV-2 synthesized DNA or extracted viral genomic-RNA were spiked into the collected nasopharyngeal and oropharyngeal swab samples. The simulated swab specimens containing the synthetic-DNA were 10-fold diluted in the nuclease-free water and 1 μl of the diluted sample was added to the iNAAT reactions. The samples containing the viral RNA were 50-fold diluted in the nuclease-free water and 5 μl of the diluted sample were added to the iNAAT reactions. Non-spiked specimens were used as negative samples.

### Preparation of lyophilized reagents for LAMP assay

A 0.2 ml reaction tube containing 7.5 μl of WarmStart Colorimetric LAMP 2X Master Mix and primers (0.1 μM each F3 and B3, 0.8 μM each FIP and BIP, and 0.2 μM each FLoop and Bloop) was mixed well and subjected to lyophilization in a freeze-dryer (Operon, South Korea) using a protocol based on the method described by Saleki-Gerhardt and Zografi [50].

## RESULTS

### Colorimetric iNAATs for SARS-CoV-2 detection

The extracted genomic RNA of SARS-CoV-2 was utilized as the template to perform the LAMP, CPA, and PSR reactions in the presence of the pH-sensitive indicator. The results indicated that the color change from red to yellow of the iNAAT reactions corresponding to the amplified products generated only when the template was present (Fig. S1, SI1 file). Next, the optimal temperature and required time of iNAAT for the detection of SARS-CoV-2 were defined. For LAMP, 30 min was the minimum time required for the readout of positive amplification judged by the naked eye (Fig. 1A, left panel) and the amplicons were produced from 60 to 70 °C (Fig. 1A, right panel). Regarding CPA and PSR, amplified products could be well visualized after 40 min (Fig. 1B and C, left panels). Meanwhile, 61 and 63 °C were the minimal temperatures for observing CPA and PSR amplified products (Fig. 1B and C, right panels). Therefore, the incubation temperatures of the iNAAT reaction were set at 60 °C for LAMP and 63 °C for CPA and PSR. The incubation time was selected as 45 min for the most significant color shift. Noted that, the color of negative control occasionally started to change to yellow only after 60 min of incubation due to the spurious amplification. This is much longer than 30 min as reported in the other study [17]. We reason that the difference resulted from the differences in the sequences and amounts of primers used. Given that the majority of false-positive results of LAMP arises from the formation of primer-dimers [17], this result indicates that the primer conditions we developed can reduce the false-positive rate of the reaction in practical usage.

**Figure 1.**
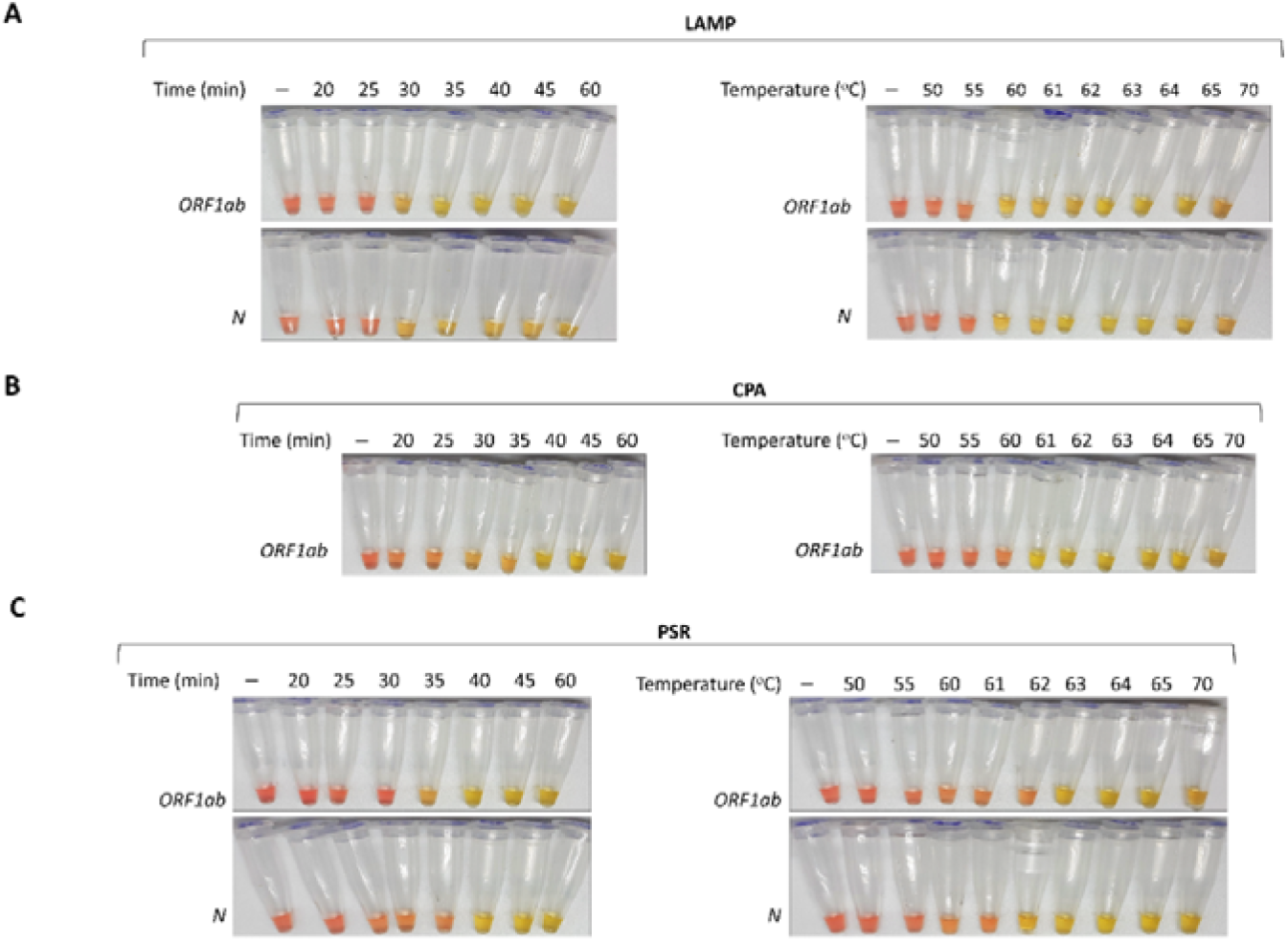
Optimization of the incubation time and temperature for iNAATs. The LAMP (in A), CPA (in B) and PSR (in C) reactions were incubated from 20 to 60 min at 60 °C (LAMP) and 63 °C (CPA and PSR) (left panels) and at different temperatures from 50 to 70 °C for 30 min (right panels).

### Specificity of SARS-CoV-2 colorimetric iNAATs

The sequences of iNAATs primers used were aligned to genome sequences of different strains of coronaviruses including 13 SARS-CoV-2 strains, MERS-CoV, SARS-CoV, human coronaviruses related to the common cold (HKU1, OC43, NL63, and 229E), bat SARS-like-CoV, Murine hepatitis virus (*Murine coronavirus*), and *Betacoronavirus England* 1. The results showed that 0% mismatch with all tested SARS-CoV-2 variants (MN938384, MN975262, MN985325, MN988668, MN988669, MN988713, MN994467, MN994468, MN997409, MT007544, MT121215, MT123292 and NC045512) was observed, suggesting that the developed iNAATs could detect different variants of SARS-CoV-2 (Table 2). In contrast, except for bat SARS-like-CoV 2015 and 2017 strains, most of the genomic RNA sequences of other coronaviruses showed nucleotide mismatch higher than 20% with the designed primers. Thus, it is likely that the designed primer sets would not amplify those sequences, ensuring the specificity of iNAATs for SARS-CoV-2. The results of *in silico* PCR and a virtual LAMP tool (Electric LAMP [51]) also supported the high specificity of primer sets used (SI2 file). Further data demonstrated that the iNAATs primer sets designed selectively detected the presence of SARS-CoV-2 DNA while no cross-reactivity was observed with SARS-CoV, MERS-CoV, bat SARS-like-CoV (Fig. 2) and some other coronaviruses (Fig. S2, SI1 file), confirming the absolute specificity of the assays.

**Figure 2.**
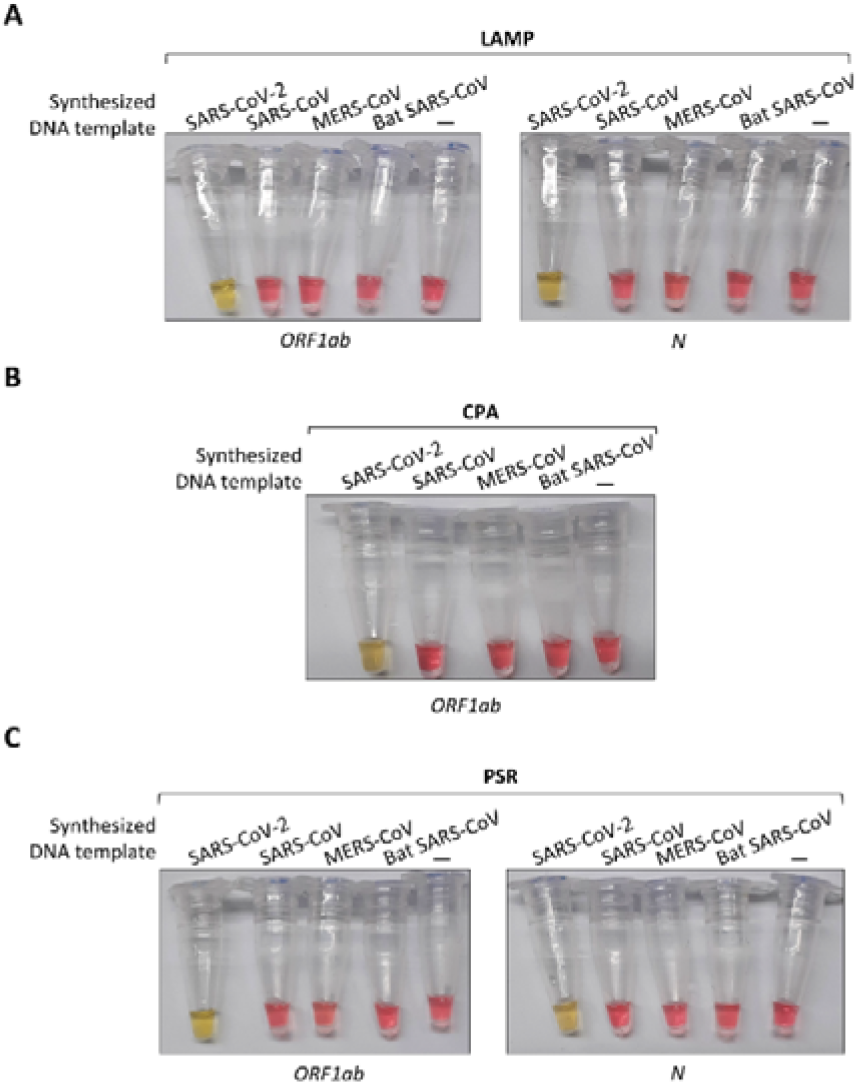
The specificity of the colorimetric iNAATs. The specificity of LAMP (in A), CPA (in B) and PSR (in C) assay was evaluated with the synthesized DNA (1 ng) of SARS-CoV-2, SARS-CoV, MERS-CoV, and bat SARS-like-CoV.

**Table 2.**
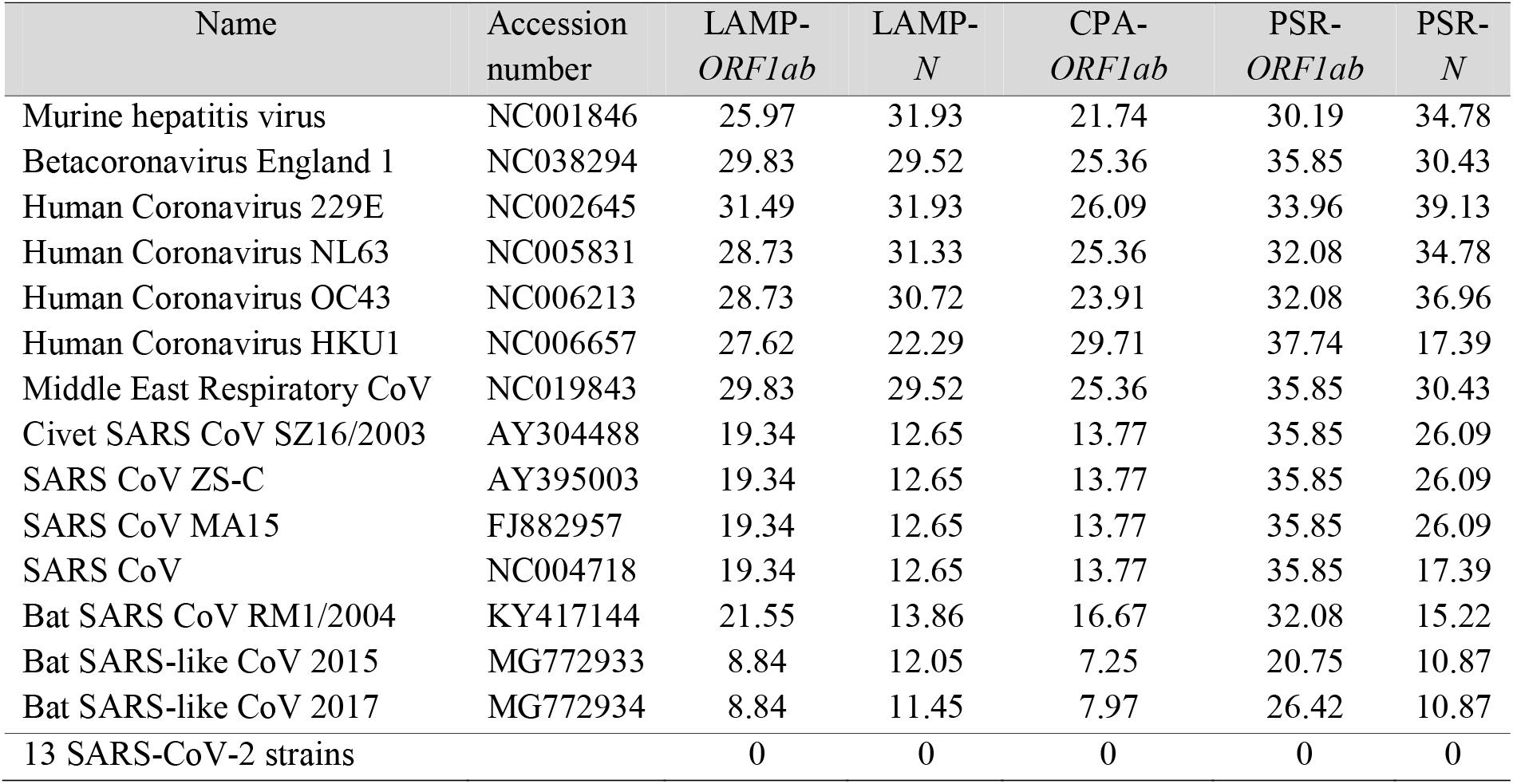
The percent mismatch of newly designed primers between SARS-CoV-2 and related taxa

| Name | Accession number | LAMP-*ORF1ab* | LAMP-*N* | CPA-*ORF1ab* | PSR-*ORF1ab* | PSR-*N* |
| --- | --- | --- | --- | --- | --- | --- |
| Murine hepatitis virus | NC001846 | 25.97 | 31.93 | 21.74 | 30.19 | 34.78 |
| Betacoronavirus England 1 | NC038294 | 29.83 | 29.52 | 25.36 | 35.85 | 30.43 |
| Human Coronavirus 229E | NC002645 | 31.49 | 31.93 | 26.09 | 33.96 | 39.13 |
| Human Coronavirus NL63 | NC005831 | 28.73 | 31.33 | 25.36 | 32.08 | 34.78 |
| Human Coronavirus OC43 | NC006213 | 28.73 | 30.72 | 23.91 | 32.08 | 36.96 |
| Human Coronavirus HKU1 | NC006657 | 27.62 | 22.29 | 29.71 | 37.74 | 17.39 |
| Middle East Respiratory CoV | NC019843 | 29.83 | 29.52 | 25.36 | 35.85 | 30.43 |
| Civet SARS CoV SZ16/2003 | AY304488 | 19.34 | 12.65 | 13.77 | 35.85 | 26.09 |
| SARS CoV ZS-C | AY395003 | 19.34 | 12.65 | 13.77 | 35.85 | 26.09 |
| SARS CoV MA15 | FJ882957 | 19.34 | 12.65 | 13.77 | 35.85 | 26.09 |
| SARS CoV | NC004718 | 19.34 | 12.65 | 13.77 | 35.85 | 17.39 |
| Bat SARS CoV RM1/2004 | KY417144 | 21.55 | 13.86 | 16.67 | 32.08 | 15.22 |
| Bat SARS-like CoV 2015 | MG772933 | 8.84 | 12.05 | 7.25 | 20.75 | 10.87 |
| Bat SARS-like CoV 2017 | MG772934 | 8.84 | 11.45 | 7.97 | 26.42 | 10.87 |
| 13 SARS-CoV-2 strains |  | 0 | 0 | 0 | 0 | 0 |

### Limit of detection of colorimetric iNAATs

The LOD values of iNAAT reactions were determined using synthesized DNA templates serially diluted in nuclease-free water. As shown in Fig. 3A and B, roughly a single copy of the synthesized targeted gene per reaction was the lowest template amount that iNAAT reaction can detect. As for PSR, the LOD of the reaction targeting the *ORF1ab* sequence was 10^3^ copies/reaction (Fig. 3C). The obtained results indicated that the iNAAT reactions containing primer pairs designed for LAMP, CPA and PSR targeting *N*-sequence were sensitive enough for the practical diagnosis.

**Figure 3.**
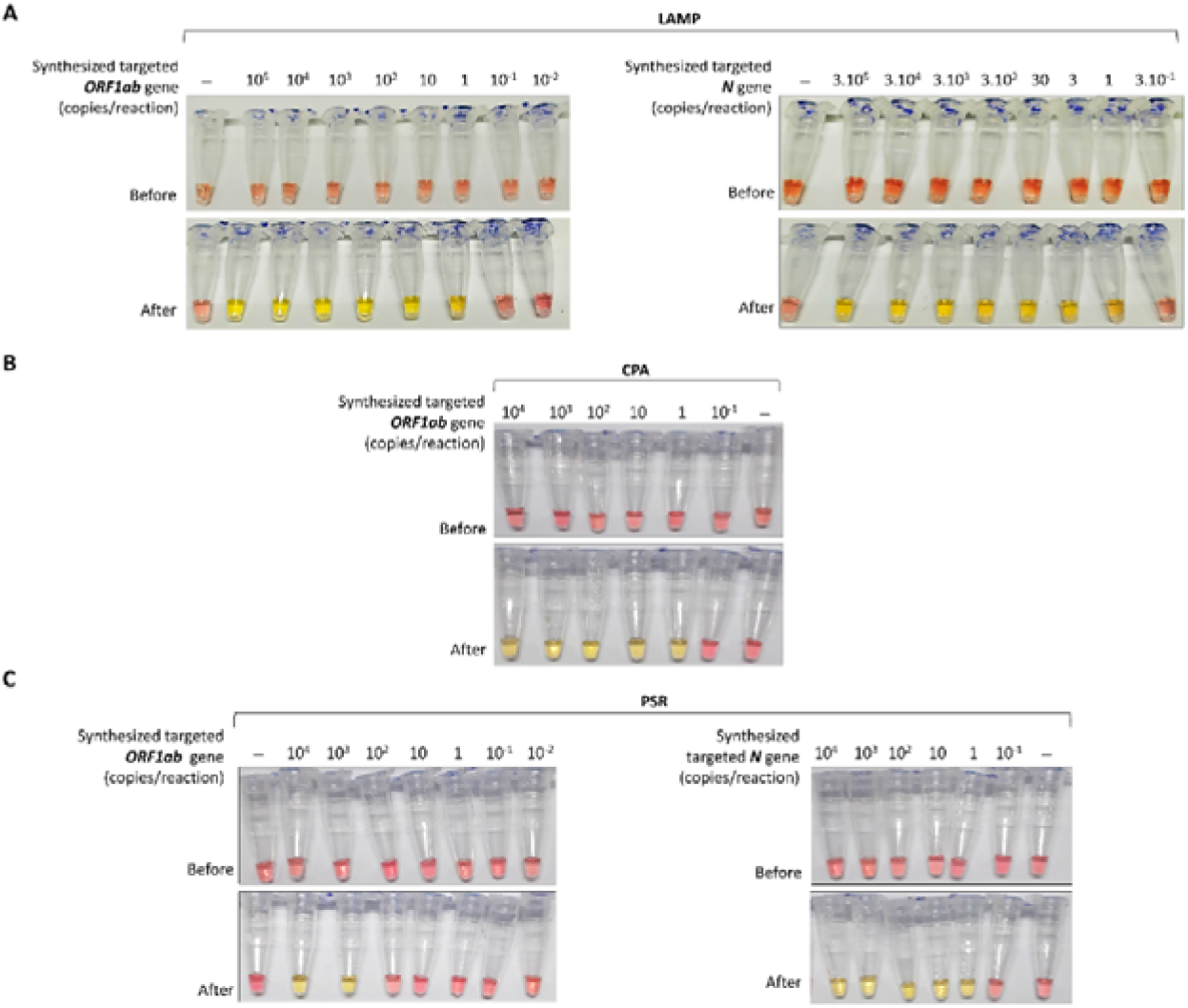
The LOD values for the targeted genes of colorimetric iNAATs. LOD of LAMP (in A), CPA (in B) and PSR (in C) assay were evaluated. The synthesized DNA template was serially diluted in nuclease-free water to the indicated concentrations and 1 μl of the diluted DNA templates was added to the reactions. The experiments were replicated at least 3 times.

The genomic RNA of SARS-CoV-2 was extracted and quantitated using the standard curve based on qRT-PCR Ct-value as described in the previous study [49]. The LODs of the colorimetric iNAATs on this extracted SARS-CoV-2 genomic RNA were also evaluated, revealing the LOD values of LAMP reactions were around 21.57 (*ORF1ab*) and 43.14 (*N*) viral-RNA copies/reaction (Table 3). The obtained values were outstanding compared to 431.47 and 862.9 genome copies/reaction of CPA and PSR assays, respectively (Table 3).

**Table 3.** LODs of colorimetric iNAATs for extracted SARS-CoV-2 genomic RNA

| Viral RNA<br>copies (per<br>reaction) | Ratio of positive tests to the total test number |  |  |  |  |  |
| --- | --- | --- | --- | --- | --- | --- |
|  | MP |  | yophilized LAMP |  | CPA | PSR |
|  | *ORF1ab* | *N* | *ORF1ab* | *N* | *ORF1ab* | *N* |
| 1725.89 | 3/3 | 3/3 | 3/3 | 3/3 | 3/3 | 3/3 |
| 862.95 | - | - | - | - | 3/3 | 3/3 |
| 431.47 | - | - | - | - | 3/3 | 1/3 |
| 215.74 | - | - | - | - | 0/3 | 0/3 |
| 172.59 | 3/3 | 3/3 | 3/3 | 3/3 | 0/3 | 0/3 |
| 86.30 | 3/3 | 3/3 | 3/3 | 3/3 | 0/3 | 0/3 |
| 43.15 | 3/3 | 3/3 | 3/3 | 3/3 | 0/3 | 0/3 |
| 21.57 | 3/3 | 0/3 | 0/3 | 0/3 | 0/3 | 0/3 |
| 17.26 | 0/3 | 0/3 | 0/3 | 0/3 | 0/3 | 0/3 |
| 1.73 | 0/3 | 0/3 | 0/3 | 0/3 | 0/3 | 0/3 |
| LOD | 21.57 | 43.14 | 43.14 | 43.14 | 431.47 | 862.95 |

### Performance of direct SARS-CoV-2 colorimetric iNAATs with simulated clinical specimens

The activity of the isothermal polymerase utilized in LAMP, CPA and PSR is more tolerant to various PCR inhibitors such as trace quantities of whole-blood, hemin, urine or stools [28, 52, 53]. Therefore, we attempted to use these iNAATs to directly detect SARS-CoV-2 nucleic acid from nasopharyngeal and oropharyngeal swab specimens. Synthesized DNAs of SARS-CoV-2 were spiked into the crude samples to mimic the clinical specimens. Unfortunately, the direct use of undiluted crude samples interfered with the color indicator of the iNAATs; the reaction color changed to yellow immediately when samples were added. As expected, sample dilution was required to clearly establish the colorimetric reactions between positive and negative signals (data not shown). Accordingly, a 10-fold dilution of nasopharyngeal and oropharyngeal swab specimens, which was sufficient to distinguish the colors between positive and negative samples, was selected to examine the simulated clinical specimens prepared.

As a result, all nasopharyngeal and oropharyngeal swab samples without spiked DNA produced negative signals while the samples containing a various number of spiked DNAs produced positive signals (Fig. 4), indicating that iNAATs successfully detected the nucleic acid of SARS-CoV-2 in crude specimens. Among the three methods, based on the best performance on detecting genomic RNA of SARS-CoV-2, LAMP assay was used for further evaluation with simulated clinical samples varied in the amounts of spiked viral-RNAs. Note that instead of transferring 1 μl of the 10-fold diluted samples to the reaction, 5 μl of the spiked viral-RNA specimens were added to the reaction. Thus, a 50-fold dilution of the simulated specimens was prepared proportionally. The results indicated that the LAMP assays can directly detect SARS-CoV-2 genomic-RNA in crude samples (Fig. 5), supporting the success of the direct SARS-CoV-2 colorimetric LAMP assay designed.

**Figure 4.**
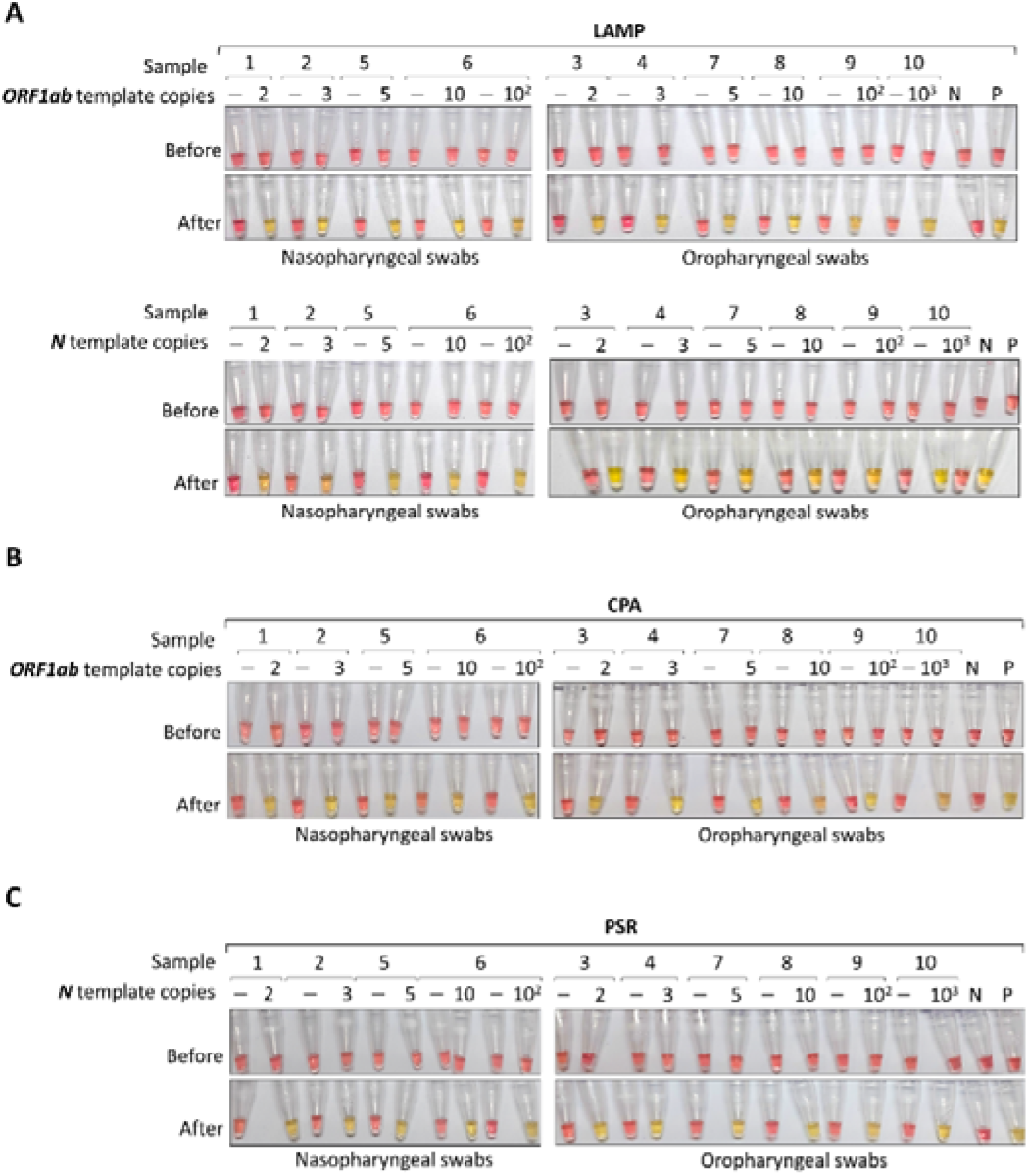
The direct iNAATs for detecting SARS-CoV-2 in simulated samples. Different amounts of the synthetic-DNA template were spiked into the nasopharyngeal and oropharyngeal swab samples to simulate the clinical samples containing SARS-CoV-2, and 1 μl of the 10-fold diluted specimens was added to the LAMP (in A), CPA (in B) and PSR (in C) reactions. The number indicates the copy of template per reaction. Abbreviation, N: negative control; P: positive control.

**Figure 5.**
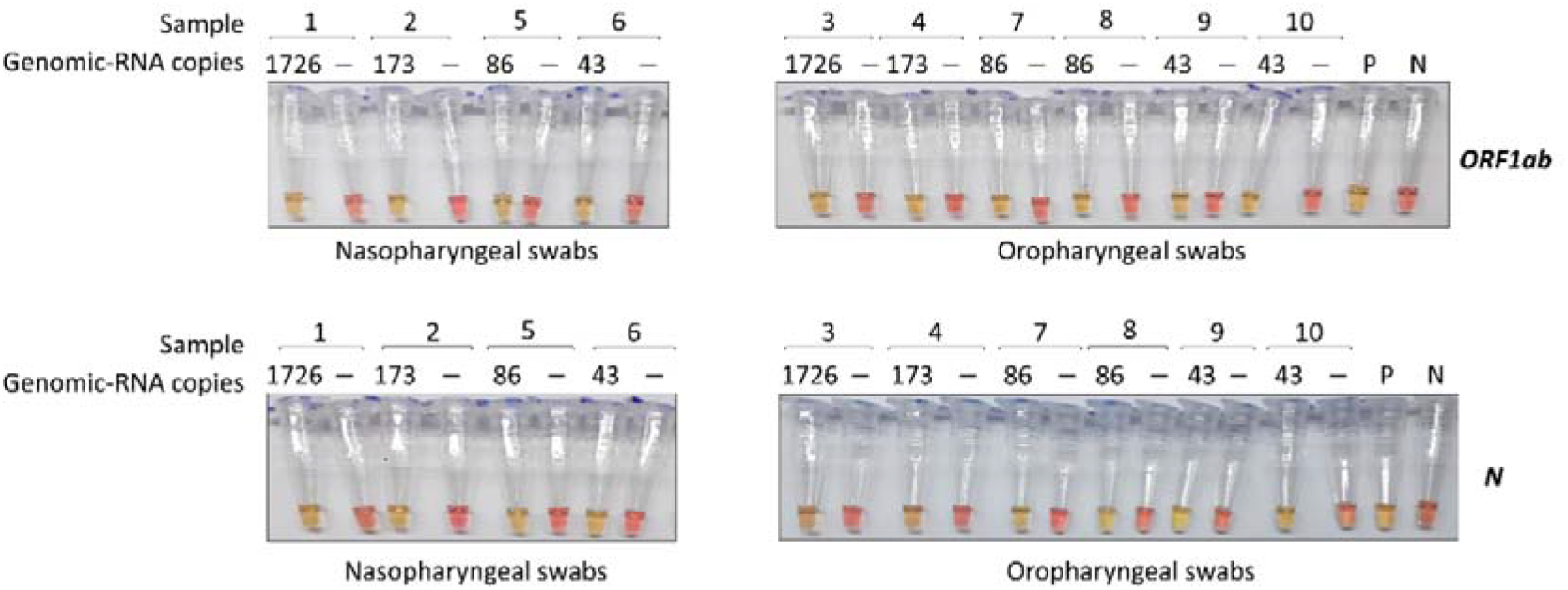
Colorimetric LAMP assays for detection of SARS-CoV-2 genomic RNA in simulated samples. Various concentrations of genomic RNA of SARS-CoV-2 were spiked into the nasopharyngeal and oropharyngeal swab samples to simulate the clinical specimens containing SARS-CoV-2. The mimicked samples were 50-fold diluted and 5 μl of the diluted specimens were added to the LAMP reactions. The number indicates the copy of viral RNA per reaction. Abbreviation, N: negative control; P: positive control.

### Performance of SARS-CoV-2 colorimetric LAMP using lyophilized reagents

For better on-site testing, lyophilized reagents that are ready-to-use without strict storage conditions at low temperature would be highly advantageous. Thus, we also evaluated the LAMP kit performance using the dried reagents. The lyophilized kit exhibited the same LOD value of 1 DNA copy/reaction, which is equivalent to 10 DNA copies/μl in simulated specimens (Fig. 6A). Subsequently, the LOD values of lyophilized LAMP kit for the detection of SARS-CoV-2 genomic RNA were also evaluated. Both primer sets could identify approximately 43.14 copies of viral RNA per reaction (Table 3). Note that the LOD values defined correspond to a Ct value of roughly 36.5 when tested by qRT-PCR for the *E* gene [49]. This high Ct value (>35) strongly indicates that the designed lyophilized LAMP kit could identify the specimens with low-level infection of SARS-CoV-2 found during early infection or asymptomatic carriage. Importantly, all mimicked samples containing different amounts of synthesized DNA or viral RNA were detected by the lyophilized LAMP kit (Fig. 6B and C). The direct addition of unextracted clinical specimens to the reaction can markedly reduce the time required for sample preparation and thus, simplify the operation procedure. When all those features are considered, the lyophilized LAMP kit is highly suitable for POC diagnosis.

**Figure 6.**
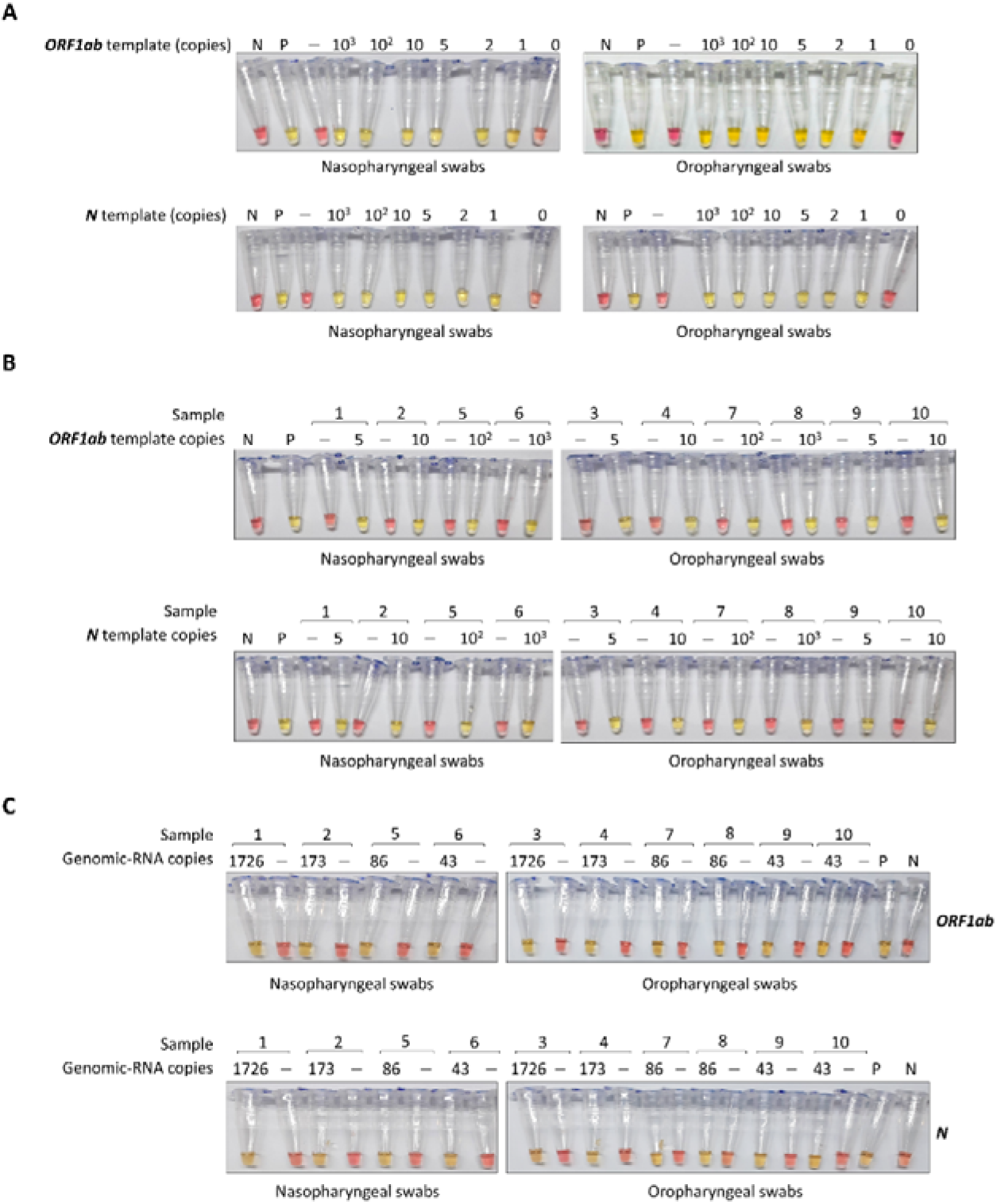
The performance of SARS-CoV-2 colorimetric lyophilized LAMP kit. (A) LODs of lyophilized LAMP kit evaluated in the simulated samples. The defined amount of synthetic DNA templates were spiked into the nasopharyngeal and oropharyngeal swab samples, followed by the 10-fold dilution of simulated specimens into water and 1 μl of the diluted samples was added to the reaction. The sensitivity of lyophilized LAMP kit was examined with mimicked clinical specimens containing different amounts of spiked synthetic DNA (in B) and SARS-CoV-2 genomic RNA (in C). The number indicates the copy number of synthetic DNA or viral RNA in reaction. Abbreviation, N: negative control; P: positive control.

## DISCUSSION

The pneumonia outbreak COVID-19 has recently become a global pandemic. While there has been not a commercial vaccine or efficient chemotherapeutics successfully developed, rapid diagnosis and necessary biosecurity procedures are the most essential actions to control the disease. POC such as airports or stations should be strictly managed to avoid spreading COVID-19. However, the prevention measures including body heat monitoring, checking personal travel history and clinical symptoms have been proven to be insufficient as many asymptomatic people are infectious [54–57]. Self-isolation and group quarantine of suspected cases can temporarily limit the transmission, but it will be more difficult to control when the number of suspects is high. While the gold standard for identifying the patients is qRT-PCR, the standard operating procedure of diagnosis including preparation of master mix, extraction of nucleic acid template and requirement of expensive qRT-PCR devices to examine multiple specimens simultaneously is not highly portable. Moreover, the testing time required is at least a few hours, if not including the time needed for transferring the sample to the labs, thus, slowing down the quarantine process of the infected patient. Consequently, the qRT-PCR tests only serve to verify a small number of cases while the actions cannot be made immediately. Therefore, POC diagnostic detection methods that are rapid and accurate can help the authority to effectively monitor the spreading of the virus.

The three isothermal amplification methods interested including LAMP, CPA and PSR all require the DNA polymerase strand displacement activity. Among them, LAMP was first introduced and has been widely utilized [25, 26, 58]. Meanwhile, CPA and PSR were later developed and studies have shown that their performance was comparable to LAMP assay [38–42]. Moreover, since the robust amplified process of LAMP, CPA and PSR techniques induces the proton release which significantly drops the pH of reaction, allowing the reaction outcome can be easily read by the naked eye with the use of an economic pH-sensitive indicator, thus simplifying the handling process and reducing the cost of assays. In contrast, detection based on the pH change is more limited for PCR or other isothermal methods such as RPA (Recombinase polymerase amplification) or HDA (Helicase-dependent amplification). In response to the COVID-19 pandemic, multiple LAMP assays were introduced and evaluated to rapidly detect SARS-CoV-2 in patient samples [12–15, 17]. In this work, for the first time, the three methods including LAMP. CPA and PSR were applied to detect SARS-CoV-2 to compare their effectiveness. Surprisingly, despite showing similar performance regarding the identification of the synthetic DNA template of SARS-CoV-2, LAMP was superior to CPA and PSR in the detection of genomic RNA of SARS-CoV-2. The finding indicates that LAMP is the best among the three isothermal techniques evaluated to be used as the diagnostic method for the identification of SARS-CoV-2 in practice. This is consistent with the fact that many recent studies have attempted to develop LAMP assays for COVID-19 diagnosis [12–15, 17].

One of the most distinctive features of the SARS-CoV-2 colorimetric LAMP kit developed herein is the direct use of minimally processed clinical samples. The only step required to prepare samples before adding to the reaction is simple dilution. The study was inspired by different previous results showing that LAMP was successfully conducted in crude clinical specimens without sample extraction [59, 60]. We speculate that the crude samples already contained some free viral RNA and that the moderately high temperature (60 °C) of the incubation process could help to break the virus envelop, thereby releasing more genomic RNA into the solution. To our knowledge, initially, this work had been one of the first few reports attempting to develop the RT-LAMP assay for SARS-CoV-2 detection using unextracted nasopharyngeal and oropharyngeal swab specimens. Recently, several works also proved that RT-LAMP can directly detect the presence of SARS-CoV-2 in unextracted clinical samples [17, 61], further corroborating our findings. Without the requirement for viral RNA extraction, the colorimetric lyophilized LAMP kit developed is very suitable for on-site diagnosis, not only because it is much less time-consuming and laborious but also does not depend on RNA extraction kits that have been in shortage due to overwhelming global demand.

In general, the colorimetric lyophilized RT-LAMP kit developed in this study possesses the following features: (i) fast detection of SARS-CoV-2 genomic-RNA directly from the nasopharyngeal and oropharyngeal specimens within 45 minutes; (ii) high sensitivity (roughly 43 copies of the viral RNA in the reaction is sufficient for detection); (iii) naked-eye readout of results; (iv) possible detection of the virus in the early stage of infection; (v) only a common thermal incubator is needed; (vi) is amenable to high throughput testing; (vii) portable to use in any place and does not require specialized personnel to do the test; and (viii) particularly useful for resource-limited settings. This RT-LAMP kit is hence more scalable for mass testing and a promising candidate for POC diagnosis of COVID-19.

The colorimetric direct RT-LAMP method established here has been developed into ready-to-use kits. Because the number of COVID-19 patients in Vietnam was limited and the clinical COVID-19 specimens were not accessible, the authors were unable to carry out clinical trials for the kit in Vietnam. Thus, we collaborated with the researchers at Universitas Indonesia in Indonesia in an attempt to validate our kits. The initial data obtained by our collaborators revealed that the sensitivity and specificity of the kits were both over 80% (personal communication). The obtained data adamantly support that the direct RT-LAMP kit developed in this study is suitable for practical use as a rapid screening method for COVID-19 patients. The positive cases can be further tested by qRT-PCR to verify the results. This result sets a solid foundation for future extensive clinical trials of our kit.

## CONCLUSION

In this study, the colorimetric iNAATs for SARS-CoV-2 detection based on three methods including LAMP, CPA, and PSR were developed and compared the testing performance. The results indicated that iNAATs could directly identify the genomic RNA of SARS-CoV-2 in unextracted patient specimens. The results can be easily observed with the naked eye within 45 minutes, without cross-reactivity to closely related coronaviruses. However, LAMP outperformed both CPA and PSR, exhibiting the best LOD value of approximately 43.14 genome copies per reaction. Additionally, the ready-to-use lyophilized reagents for LAMP reactions maintained the testing performance of the liquid assays, showing that the colorimetric lyophilized LAMP kit developed for detecting SARS-CoV-2 nucleic acids is feasible for POC diagnosis settings.

## Supporting information

In silico PCR and eLAMP

## Data Availability

The authors confirm that the data supporting the findings of this study are available within the article and its supplementary materials and are available from the corresponding author upon request.

## Ethical statement

The sample collection was approved by Hospital Management. The internal use of samples was agreed upon under the medical and ethical rules of each participating individuals. The study was approved by the Research Ethics Committee of Pasteur Institute in Ho Chi Minh City, Vietnam.

## Acknowledgements

The authors are grateful to Professor Le Van Phan at College of Veterinary Medicine, Vietnam National University of Agriculture, Hanoi, Vietnam for sharing genomic RNAs of some coronaviruses. The authors are also thankful to the staff of the local hospital for collecting samples and providing the epidemiological information.

## Supplementary materials

SI1 file presents the results of colorimetric iNAATs to detect the presence of SARS-CoV-2 RNA and the evaluation of specificity of the colorimetric iNAATs regarding some other coronaviruses. SI2 file shows the results of *in silico* PCR and eLAMP.

## REFERENCES

1. Gaunt ER, Hardie A, Claas EC, Simmonds P, Templeton KE: Epidemiology and clinical presentations of the four human coronaviruses 229E, HKU1, NL63, and OC43 detected over 3 years using a novel multiplex real-time PCR method. Journal of clinical microbiology 2010, 48:2940–2947.

2. Zeng Z-Q, Chen D-H, Tan W-P, Qiu S-Y, Xu D, Liang H-X, Chen M-X, Li X, Lin Z-S, Liu W-K: Epidemiology and clinical characteristics of human coronaviruses OC43, 229E, NL63, and HKU1: a study of hospitalized children with acute respiratory tract infection in Guangzhou, China. European Journal of Clinical Microbiology & Infectious Diseases 2018, 37:363–369.

3. World Health Organization: Consensus document on the epidemiology of severe acute respiratory syndrome (SARS). World Health Organization; 2003.

4. Zumla A, Hui DS, Perlman S: Middle East respiratory syndrome. The Lancet 2015, 386:995–1007.

5. World Health Organization: Surveillance case definitions for human infection with novel coronavirus (nCoV): interim guidance v1, January 2020. World Health Organization; 2020.

6. Cucinotta D, Vanelli M: WHO declares COVID-19 a pandemic. Acta bio-medica: Atenei Parmensis 2020, 91:157–160.

7. COVID-19 CORONAVIRUS PANDEMIC [https://www.worldometers.info/coronavirus/]

8. He F, Deng Y, Li W: Coronavirus disease 2019: What we know? Journal of medical virology 2020, 92:719–725.

9. Naqvi AAT, Fatima K, Mohammad T, Fatima U, Singh IK, Singh A, Atif SM, Hariprasad G, Hasan GM, Hassan MI: Insights into SARS-CoV-2 genome, structure, evolution, pathogenesis and therapies: Structural genomics approach. Biochimica et Biophysica Acta (BBA)-Molecular Basis of Disease 2020:165878.

10. Yoshimoto FK: The Proteins of Severe Acute Respiratory Syndrome Coronavirus-2 (SARS CoV-2 or n-COV19), the Cause of COVID-19. The Protein Journal 2020:1.

11. World Health Organization: Novel coronavirus (2019-nCoV) technical guidance: laboratory testing for 2019-nCoV in humans. World Health Organization, Geneva, Switzerland https://www.who.int/emergencies/diseases/novel-coronavirus-2019/technical-guidance/laboratory-guidance 2020.

12. Park G-S, Ku K, Baek S-H, Kim S-J, Kim SI, Kim B-T, Maeng J-S: Development of Reverse Transcription Loop-mediated Isothermal Amplification (RT-LAMP) Assays Targeting SARS-CoV-2. The Journal of Molecular Diagnostics 2020.

13. Huang W, Boon L, Hsu C, Xiong D, Wu W, Yu Y, Zeng Y, Jia H, Zhang X, Chang H: RT-LAMP for rapid diagnosis of coronavirus SARS-CoV-2. Microbial Biotechnology 2020.

14. Lu R, Wu X, Wan Z, Li Y, Jin X, Zhang C: A Novel Reverse Transcription Loop-Mediated Isothermal Amplification Method for Rapid Detection of SARS-CoV-2. International Journal of Molecular Sciences 2020, 21:2826.

15. Yan C, Cui J, Huang L, Du B, Chen L, Xue G, Li S, Zhang W, Zhao L, Sun Y: Rapid and visual detection of 2019 novel coronavirus (SARS-CoV-2) by a reverse transcription loop-mediated isothermal amplification assay. Clinical Microbiology and Infection 2020.

16. Behrmann O, Bachmann I, Spiegel M, Schramm M, El Wahed AA, Dobler G, Dame G, Hufert FT: Rapid detection of SARS-CoV-2 by low volume real-time single tube reverse transcription recombinase polymerase amplification using an exo probe with an internally linked quencher (exoIQ). Clinical Chemistry 2020.

17. Thi VLD, Herbst K, Boerner K, Meurer M, Kremer LP, Kirrmaier D, Freistaedter A, Papagiannidis D, Galmozzi C, Stanifer ML: A colorimetric RT-LAMP assay and LAMP-sequencing for detecting SARS-CoV-2 RNA in clinical samples. Science Translational Medicine 2020, 12.

18. Broughton JP, Deng X, Yu G, Fasching CL, Servellita V, Singh J, Miao X, Streithorst JA, Granados A, Sotomayor-Gonzalez A: CRISPR–Cas12-based detection of SARS-CoV-2. Nature Biotechnology 2020:1–5.

19. Liu W, Liu L, Kou G, Zheng Y, Ding Y, Ni W, Wang Q, Tan L, Wu W, Tang S: Evaluation of Nucleocapsid and Spike Protein-based ELISAs for detecting antibodies against SARS-CoV-2. Journal of Clinical Microbiology 2020.

20. Okba N, Müller MA, Li W, Wang C, GeurtsvanKessel CH, Corman VM, Lamers MM, Sikkema RS, de Bruin E, Chandler FD: Severe Acute Respiratory Syndrome Coronavirus 2-Specific Antibody Responses in Coronavirus Disease 2019 Patients. Emerging infectious diseases 2020, 26.

21. Deng J, Jin Y, Liu Y, Sun J, Hao L, Bai J, Huang T, Lin D, Jin Y, Tian K: Serological survey of SARS□CoV□2 for experimental, domestic, companion and wild animals excludes intermediate hosts of 35 different species of animals. Transboundary and Emerging Diseases 2020.

22. Stadlbauer D, Amanat F, Chromikova V, Jiang K, Strohmeier S, Arunkumar GA, Tan J, Bhavsar D, Capuano C, Kirkpatrick E: SARS□CoV□2 Seroconversion in Humans: A Detailed Protocol for a Serological Assay, Antigen Production, and Test Setup. Current Protocols in Microbiology 2020, 57:e100.

23. Yan Y, Chang L, Wang L: Laboratory testing of SARS□CoV, MERS□CoV, and SARS□CoV□2 (2019□nCoV): Current status, challenges, and countermeasures. Reviews in Medical Virology 2020:e2106.

24. Notomi T, Okayama H, Masubuchi H, Yonekawa T, Watanabe K, Amino N, Hase T: Loop-mediated isothermal amplification of DNA. Nucleic Acids Res 2000, 28:E63.

25. Mori Y, Notomi T: Loop-mediated isothermal amplification (LAMP): Expansion of its practical application as a tool to achieve universal health coverage. Journal of Infection and Chemotherapy 2020, 26:13–17.

26. Uwiringiyeyezu T, El Khalfi B, Belhachmi J, Soukri A: Loop-mediated Isothermal Amplification LAMP, Simple Alternative Technique of Molecular Diagnosis Process in Medicals Analysis: A Review. Annual Research & Review in Biology 2019:1–12.

27. Becherer L, Borst N, Bakheit M, Frischmann S, Zengerle R, von Stetten F: Loop-mediated isothermal amplification (LAMP)–review and classification of methods for sequence-specific detection. Analytical Methods 2020, 12:717–746.

28. Dhama K, Karthik K, Chakraborty S, Tiwari R, Kapoor S, Kumar A, Thomas P: Loop-mediated isothermal amplification of DNA (LAMP): a new diagnostic tool lights the world of diagnosis of animal and human pathogens: a review. Pak J Biol Sci 2014, 17:151–166.

29. Zhang X, Lowe SB, Gooding JJ: Brief review of monitoring methods for loop-mediated isothermal amplification (LAMP). Biosensors and Bioelectronics 2014, 61:491–499.

30. Lin Z, Zhang Y, Zhang H, Zhou Y, Cao J, Zhou J: Comparison of loop-mediated isothermal amplification (LAMP) and real-time PCR method targeting a 529-bp repeat element for diagnosis of toxoplasmosis. Veterinary parasitology 2012, 185:296–300.

31. Bista BR, Ishwad C, Wadowsky RM, Manna P, Randhawa PS, Gupta G, Adhikari M, Tyagi R, Gasper G, Vats A: Development of a loop-mediated isothermal amplification assay for rapid detection of BK virus. Journal of clinical microbiology 2007, 45:1581–1587.

32. Sugiyama H, Yoshikawa T, Ihira M, Enomoto Y, Kawana T, Asano Y: Comparison of loop? mediated isothermal amplification, real time PCR, and virus isolation for the detection of herpes simplex virus in genital lesions. Journal of medical virology 2005, 75:583–587.

33. Parida M, Shukla J, Sharma S, Rao PL: Rapid and real-time detection of human viral infections: current trends and future perspectives. Proceedings of the National Academy of Sciences, India Section B: Biological Sciences 2012, 82:199–207.

34. Li C, Li Z, Jia H, Yan J: One-step ultrasensitive detection of microRNAs with loop-mediated isothermal amplification (LAMP). Chemical Communications 2011, 47:2595–2597.

35. Fang R, Li X, Hu L, You Q, Li J, Wu J, Xu P, Zhong H, Luo Y, Mei J: Cross-priming amplification for rapid detection of Mycobacterium tuberculosis in sputum specimens. J Clin Microbiol 2009, 47:845–847.

36. Xu G, Hu L, Zhong H, Wang H, Yusa S-i, Weiss TC, Romaniuk PJ, Pickerill S, You Q: Cross priming amplification: mechanism and optimization for isothermal DNA amplification. Scientific reports 2012, 2:246.

37. Liu W, Dong D, Yang Z, Zou D, Chen Z, Yuan J, Huang L: Polymerase spiral reaction (PSR): a novel isothermal nucleic acid amplification method. Scientific reports 2015, 5:12723.

38. Niczyporuk JS, Woźniakowski G, Samorek-Salamonowicz E: Application of crosspriming amplification (CPA) for detection of fowl adenovirus (FAdV) strains. Archives of virology 2015, 160:1005–1013.

39. Woźniakowski G, Frączyk M, Kowalczyk A, Pomorska-Mól M, Niemczuk K, Pejsak Z: Polymerase cross-linking spiral reaction (PCLSR) for detection of African swine fever virus (ASFV) in pigs and wild boars. Scientific reports 2017, 7:1–10.

40. Ji J, Xu X, Wang X, Zuo K, Li Z, Leng C, Kan Y, Yao L, Bi Y: Novel polymerase spiral reaction assay for the visible molecular detection of porcine circovirus type 3. BMC veterinary research 2019, 15:322.

41. Sun W, Du Y, Li X, Du B: Rapid and Sensitive Detection of Hepatitis C Virus in Clinical Blood Samples Using Reverse Transcriptase Polymerase Spiral Reaction. Journal of Microbiology and Biotechnology 2020, 30:459–468.

42. Zhao L, Niu J, Gao X, Liu C, Liu S, Jiang N, Lv X, Zheng S: Development and application of isothermal amplification methods for rapid detection of F4 fimbriae producing Escherichia coli. Polish Journal of Veterinary Sciences 2020:143–152.

43. Tanner NA, Zhang Y, Evans TC, Jr.: Visual detection of isothermal nucleic acid amplification using pH-sensitive dyes. Biotechniques 2015, 58:59–68.

44. COVID-19 TREATMENT AND VACCINE TRACKER [https://covid-19tracker.milkeninstitute.org/]

45. Wire B: Pfizer and BioNTech Announce Vaccine Candidate Against COVID-19 Achieved Success in First Interim Analysis from Phase 3 Study. 2020.

46. Gallagher J: Moderna: COVID vaccine shows nearly 95% protection. 2020b. Moderna: Covid vaccine shows nearly, 95.

47. Knoll MD, Wonodi C: Oxford–AstraZeneca COVID-19 vaccine efficacy. The Lancet 2021, 397:72–74.

48. Dyer O: Covid-19: Many poor countries will see almost no vaccine next year, aid groups warn. BMJ: British Medical Journal (Online) 2020, 371.

49. Lan PT, Cuong HQ, Linh HT, Hieu NT, Anh NH, Ton T, Dong TC, Thao VT, Tuoi DTH, Tuan ND, et al: Development of standardized specimens with known concentrations for severe acute respiratory syndrome coronavirus 2 Realtime-RT-PCR testing validation. Bull World Health Organ Epub: 20 April 2020 2020.

50. Saleki-Gerhardt A, Zografi G: Non-isothermal and isothermal crystallization of sucrose from the amorphous state. Pharmaceutical Research 1994, 11:1166–1173.

51. Salinas NR, Little DP: Electric LAMP: virtual loop-mediated isothermal AMPlification. International Scholarly Research Notices 2012, 2012.

52. Kaneko H, Kawana T, Fukushima E, Suzutani T: Tolerance of loop-mediated isothermal amplification to a culture medium and biological substances. Journal of biochemical and biophysical methods 2007, 70:499–501.

53. Francois P, Tangomo M, Hibbs J, Bonetti E-J, Boehme CC, Notomi T, Perkins MD, Schrenzel J: Robustness of a loop-mediated isothermal amplification reaction for diagnostic applications. FEMS Immunology & Medical Microbiology 2011, 62:41–48.

54. Yu X, Yang R: COVID□19 transmission through asymptomatic carriers is a challenge to containment. Influenza and Other Respiratory Viruses 2020.

55. Gandhi M, Yokoe DS, Havlir DV: Asymptomatic transmission, the Achilles’ heel of current strategies to control COVID-19. Mass Medical Soc; 2020.

56. Ye F, Xu S, Rong Z, Xu R, Liu X, Deng P, Liu H, Xu X: Delivery of infection from asymptomatic carriers of COVID-19 in a familial cluster. International Journal of Infectious Diseases 2020.

57. Al-Tawfiq JA: Asymptomatic coronavirus infection: MERS-CoV and SARS-CoV-2 (COVID-19). Travel Med Infect Dis 2020, 101608.

58. Becherer L, Borst N, Bakheit M, Frischmann S, Zengerle R, von Stetten F: Loop-mediated isothermal amplification (LAMP)–review and classification of methods for sequence-specific detection. Analytical Methods 2020.

59. Iwamoto T, Sonobe T, Hayashi K: Loop-mediated isothermal amplification for direct detection of Mycobacterium tuberculosis complex, M. avium, and M. intracellulare in sputum samples. Journal of clinical microbiology 2003, 41:2616–2622.

60. Mikita K, Maeda T, Yoshikawa S, Ono T, Miyahira Y, Kawana A: The Direct Boil-LAMP method: a simple and rapid diagnostic method for cutaneous leishmaniasis. Parasitology international 2014, 63:785–789.

61. Lalli MA, Chen X, Langmade SJ, Fronick CC, Sawyer CS, Burcea LC, Fulton RS, Heinz M, Buchser WJ, Head RD: Rapid and extraction-free detection of SARS-CoV-2 from saliva with colorimetric LAMP. medRxiv 2020.

