## Supplementary Figure for "A comparative study of isothermal nucleic acid amplification methods for SARS-CoV-2 detection at point-of-care"

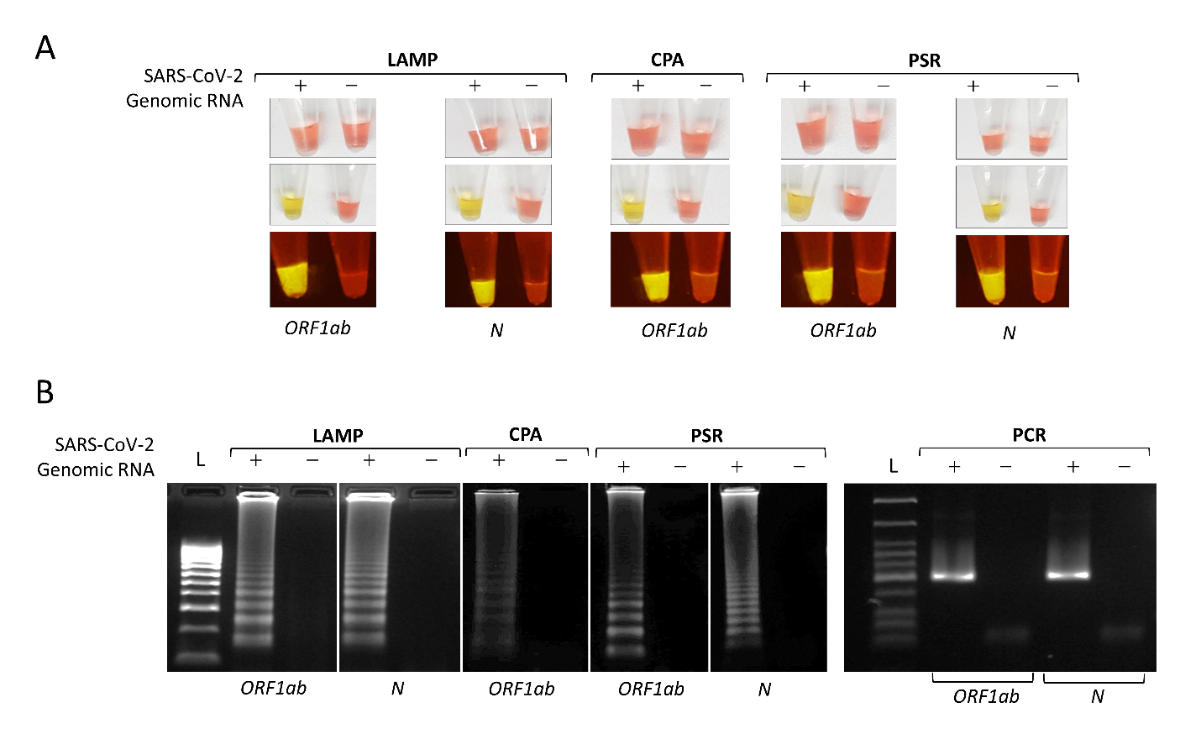

**Fig. S1.** **Colorimetric iNAATs to detect the presence of SARS-CoV-2 RNA.** (A) The color of iNAAT mixtures changed from red to yellow after incubation in the presence of SARS-CoV-2 genomic-RNA. The lowest panel: the reactions were added with SyBr Green I (Invitrogen, California, USA) and illuminated by a 470 nm light source. The reactions were incubated at 60 °C (for LAMP) and 63 °C (for CPA and PSR) for 45 min. (B) The amplified products in (A) were analyzed with gel electrophoresis and compared with the routine PCR result. The PCR assays were carried out in a 20 μl reaction volume containing 0.2 μM each of primers F3 and B3 (LAMP), 1 μl of DNA template, 0.2 µl of MyTaq DNA polymerase (Bioline, London, UK) and 4 µl of 5X MyTaq reaction buffer (Bioline, London, UK). The amplification products were analyzed by electrophoresis using a 2% agarose gel. Abbreviation, L: DNA ladder.

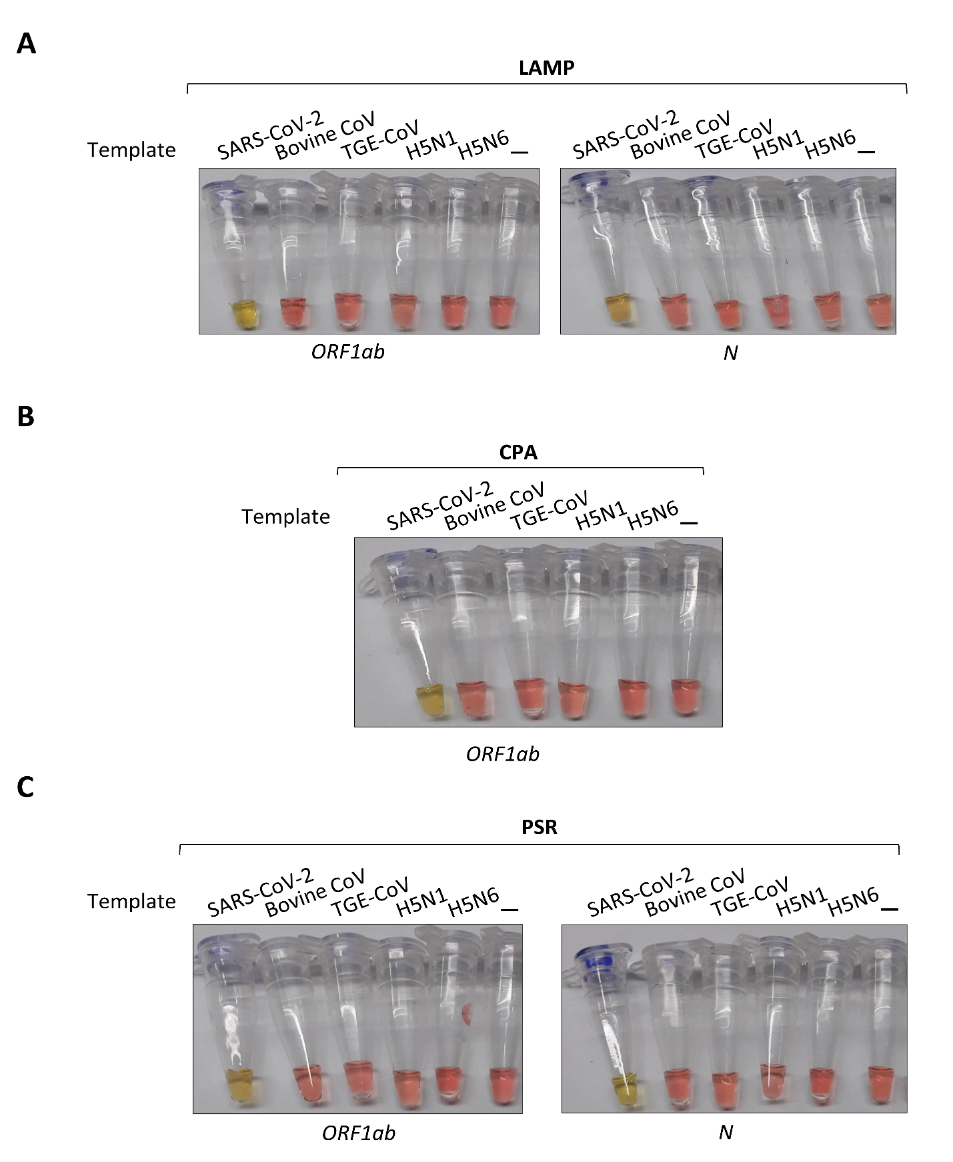

**Fig. S2.** **The specificity of the colorimetric iNAATs**. The specificity of LAMP (in A), CPA (in B) and PSR (in C) assay was evaluated with the synthesized DNA (1 ng) of SARS-CoV-2 and genomic RNAs of Bovine-CoV [1], Transmissible Gastroenteritis (TGE)-CoV in pigs, and avian influenza viruses (H5N1 and H5N6) [2].

**References**

1.	Shin, J., Tark, D., Choe, S., Cha, R. M., Park, G.-N., Cho, I.-S., Nga, B. T. T., Lan, N. T.; An, D.-J., Genetic characterization of bovine coronavirus in Vietnam. *Virus genes*, **55, (3)**, 415-420 (**2019**).

2.	Le, T. B., Kim, H. K., Na, W., Le, V. P., Song, M.-S., Song, D., Jeong, D. G.; Yoon, S.-W., Development of a multiplex RT-qPCR for the detection of different clades of avian influenza in poultry. *Viruses*, **12, (1)**, 100 (**2020**).
