## Supplementary material for "A comparative study of isothermal nucleic acid amplification methods for SARS-CoV-2 detection at point-of-care": In silico PCR and eLAMP

| **In silico PCR and eLAMP results** | | | | | | | | |
| --- | --- | --- | --- | --- | --- | --- | --- | --- |
| **In silico PCR each primer pair** | | | | | | | | |
| **0 mismatch allowed in 3'-end** | | | | | | | | |
| **Template** | **Method** | **Primer** | **Primer binding** | | | **PCR product formation** | | |
|  |  |  | **SARS-CoV-2** | **Other coronaviruses** | **Other organisms** | **SARS-CoV-2** | **Other coronaviruses** | **Other organisms** |
| ORF1ab | LAMP | FIP/BIP | + | − | − | **+** | − | − |
|  |  | F3/B3 | + | − | − | **+** | − | − |
|  |  | LoopF/LoopR | + | − | − | **−** | − | − |
|  | CPA | 4S/5A | + | − | − | **+** | − | − |
|  | PSR | F/R | + | − | − | **+** | − | − |
| N | LAMP | FIP/BiIP | + | − | − | **+** | − | − |
|  |  | F3/B3 | + | − | − | **+** | − | − |
|  |  | LoopF/LoopR | + | + (SARS) | − | **−** | − | − |
|  | PSR | F/R | + | − | − | **+** | − | − |
| **2 mismatches allowed in 3'-end** | | | | | | | | |
| **Template** | **Method** | **Primer** | **Primer binding** | | | **PCR product formation** | | |
|  |  |  | **SARS-CoV-2** | **Other coronaviruses** | **Other organisms** | **SARS-CoV-2** | **Other coronaviruses** | **Other organisms** |
| ORF1ab | LAMP | FIP/BIP | + | − | + | + | − | − |
|  |  | F3/B3 | + | − | − | + | − | − |
|  |  | LoopF/LoopR | + | − | + | − | − | + |
|  | CPA | 4S/5A | + | − | − | + | − | − |
|  | PSR | F/R | + | + (OC43) | + | + | − | + (*Staphylococcus epidermidis*) |
| N | LAMP | FIP/BiIP | + | + (SARS) | + | + | + (SARS) | − |
|  |  | F3/B3 | + | + (SARS) | + | + | − | − |
|  |  | LoopF/LoopR | + | + (SARS, 229E) | + | − | − | − |
|  | PSR | F/R | + | + (SARS, 229E) | + | + | − | − |

| **In silico PCR (each primer set)** | | | | | | | |
| --- | --- | --- | --- | --- | --- | --- | --- |
| **0 mismatch allowed in 3'-end** | | | | | | | |
| **Template** | **Method** | **Primer binding** | | | **PCR products formation** | | |
|  |  | **SARS-CoV-2** | **Other coronaviruses** | **Other organisms** | **SARS-CoV-2** | **Other coronaviruses** | **Other organisms** |
| ORF1ab | LAMP | + | − | − | + | − | − |
|  | PSR | + | − | − | + | − | − |
|  | CPA | + | − | − | + | − | − |
| N | LAMP | + | − | − | + | − | − |
|  | PSR | + | − | − | + | − | − |
| **2 mismatches allowed in 3'-end** | | | | | | | |
| **Template** | **Method** | **Primer binding** | | | **PCR products formation** | | |
|  |  | **SARS-CoV-2** | **Other coronaviruses** | **Other organisms** | **SARS-CoV-2** | **Other coronaviruses** | **Other organisms** |
| ORF1ab | LAMP | + | − | + | + | − | + (*Chlamydia psittaci, Legionella pneumophila, Mycoplasma pneumoniae, Staphylococcus epidermidis, Streptococcus pyogenes*)^1^ |
|  | PSR | + | + (OC43) | + | + | − | + (*Staphylococcus epidermidis*)^2^ |
|  | CPA | + | + (SARS, 229E) | + | + | − | − |
| N | LAMP | + | + (SARS, 229E) | + | + | + (SARS)^3^ | − |
|  | PSR | + | + (SARS, 229E) | + | + | − | − |

^1^Predicted PCR product length: *Chlamydia psittaci*: 6800 bp; *Legionella pneumophila*: 2043bp; *Mycoplasma pneumoniae*: 9589bp; *Staphylococcus epidermidis*: 7560bp; *Streptococcus pyogenes*: 737bp --> The PCR products are so long that they are hardly generated in reality

^2^*Staphylococcus epidermidis*: 3599bp

^3^SARS: 198bp (formed by FIP and BIP), and 206bp (formed by FIP and B3). F3 could not bind to SARS genome (even 5 mismatches are allowed), inhibiting the formation of loop product which is indispensable for LAMP amplification process

| List of related coronoviruses and other organisms needed to be tested for cross-reactivity that is recommended by WHO | | | | | | |
| --- | --- | --- | --- | --- | --- | --- |
|  |  | **In silico PCR (0 mismatch)** | | | | |
|  |  | LAMP-Orf1ab | LAMP-N | PSR-Orf1ab | PSR-N | CPA-Orf1ab |
|  | **Other high priority pathogens from the same virus family** |  | | | | |
|  | Human coronavirus 229E | − | − | − | − | − |
|  | Human coronavirus OC43 | − | − | − | − | − |
|  | Human coronavirus HKU1 | − | − | − | − | − |
|  | Human coronavirus NL63 | − | − | − | − | − |
|  | SARS-coronavirus | − | − | − | − | − |
|  | MERS-coronavirus | − | − | − | − | − |
|  | **High priority organisms likely in the circulating area** |  | | | | |
|  | Adenovirus | − | − | − | − | − |
|  | Human Metapneumovirus (hMPV) | − | − | − | − | − |
|  | Parainfluenza virus 1-4 | − | − | − | − | − |
|  | Influenza A | − | − | − | − | − |
|  | Influenza B | − | − | − | − | − |
|  | Enterovirus | − | − | − | − | − |
|  | Respiratory syncytial virus | − | − | − | − | − |
|  | *Rhinovirus* | − | − | − | − | − |
|  | *Chlamydia pneumonia* | − | − | − | − | − |
|  | *Legionella pneumophila* | − | − | − | − | − |
|  | *Mycobacterium tuberculosis* | − | − | − | − | − |
|  | *Streptococus pneumonia* | − | − | − | − | − |
|  | *Streptococus pyrogenes* | − | − | − | − | − |
|  | *Bordetella pertussis* | − | − | − | − | − |
|  | *Mycoplasma pneumoniae* | − | − | − | − | − |
|  | *Pneumocystis jirovecii* | − | − | − | − | − |
|  | Influenza C | − | − | − | − | − |
|  | *Parechovirus* | − | − | − | − | − |
|  | *Candida albicans* | − | − | − | − | − |
|  | *Corynebacterium diphtheriae* | − | − | − | − | − |
|  | *Legionella non pneumophila* | − | − | − | − | − |
|  | *Bacillus anthracosis* | − | − | − | − | − |
|  | *Moraxella cararrhalis* | − | − | − | − | − |
|  | *Neisseria elongate and miningitidis* | − | − | − | − | − |
|  | *Pseudomonas aeruginosa* | − | − | − | − | − |
|  | *Staphylococcus epidermis* | − | − | − | − | − |
|  | *staphylococcus salivarius* | − | − | − | − | − |
|  | *Leptospirosis* | − | − | − | − | − |
|  | *Chlamydia paittaci* | − | − | − | − | − |
|  | *Coxiella burneti* | − | − | − | − | − |
|  | *Streptococcus aureus* | − | − | − | − | − |

(-): no PCR product formed

| **eLAMP (Ref. 51)** | | |
| --- | --- | --- |
| **Tested strain** | **ORF1ab primer set** | **N primer set** |
| AE014074.1 Streptococcus pyogenes MGAS315; complete genome | 0 | 0 |
| AE017334.2 Bacillus anthracis str. 'Ames Ancestor'; complete genome | 0 | 0 |
| AJJF01000006.1 Candida albicans SC5314 supercont4.6; whole genome shotgun sequence | 0 | 0 |
| AP017922.1 Staphylococcus aureus DNA; complete genome; strain: JP080 | 0 | 0 |
| AP018036.1 Mycobacterium tuberculosis DNA; complete genome; strain: HN-506 | 0 | 0 |
| CP001019.1 Coxiella burnetii CbuG_Q212; complete genome | 0 | 0 |
| CP011448.1 Bordetella pertussis strain B3921; complete genome | 0 | 0 |
| CP015283.1 Streptococcus salivarius strain ATCC 25975; complete genome | 0 | 0 |
| CP015344.1 Legionella pneumophila strain D-7630; complete genome | 0 | 0 |
| CP025256.1 Streptococcus pneumoniae Xen35; complete genome | 0 | 0 |
| CP039256.1 Leptospira interrogans strain FMAS_KW2 chromosome I; complete sequence | 0 | 0 |
| JQ902010.1 Human parainfluenza virus 1 strain HPIV1/WI/629-D02071/2010; partial genome | 0 | 0 |
| KF268207.1 Human adenovirus 71 strain human/DEU/HEIM_00085/1987/71[P9H20F71]; complete genome | 0 | 0 |
| KJ627437.1 Human metapneumovirus strain HMPV/Homo sapiens/PER/FPP00726/2011/A; complete genome | 0 | 0 |
| KT835408.1 Enterovirus D68 strain EV68/Ontario/C818712/2014; partial genome | 0 | 0 |
| KY417148.1:28121-28362 Bat SARS-like coronavirus isolate Rs4247; complete genome | 0 | 0 |
| MN306053.1 Human coronavirus OC43 strain HCoV_OC43/Seattle/USA/SC9430/2018; complete genome | 0 | 0 |
| MN749157.1 Human rhinovirus 1B strain 12O3; complete genome | 0 | 0 |
| MN781633.1 Human parechovirus 3 isolate US-WI-09-R7; complete genome | 0 | 0 |
| MN908947.3 Severe acute respiratory syndrome coronavirus 2 isolate Wuhan-Hu-1; complete genome | 1 | 1 |
| NC_001796.2 Human parainfluenza virus 3; complete genome | 0 | 0 |
| NC_001803.1 Respiratory syncytial virus; complete genome | 0 | 0 |
| NC_002204.1 Influenza B virus RNA 1; complete sequence | 0 | 0 |
| NC_002516.2 Pseudomonas aeruginosa PAO1; complete genome | 0 | 0 |
| NC_002645.1 Human coronavirus 229E; complete genome | 0 | 0 |
| NC_003112.2 Neisseria meningitidis MC58; complete sequence | 0 | 0 |
| NC_004461.1 Staphylococcus epidermidis ATCC 12228; complete sequence | 0 | 0 |
| NC_004718.3 SARS coronavirus; complete genome | 0 | 0 |
| NC_005043.1 Chlamydia pneumoniae TW-183; complete sequence | 0 | 0 |
| NC_005831.2 Human Coronavirus NL63; complete genome | 0 | 0 |
| NC_006307.2 Influenza C virus (C/Ann Arbor/1/50) PB2 gene for polymerase 2; complete cds | 0 | 0 |
| NC_006577.2:206-21753 Human coronavirus HKU1; complete genome | 0 | 0 |
| NC_014147.1 Moraxella catarrhalis BBH18; complete genome | 0 | 0 |
| NC_017287.1 Chlamydia psittaci 6BC; complete sequence | 0 | 0 |
| NC_019843.3 Middle East respiratory syndrome-related coronavirus isolate HCoV-EMC/2012; complete genome | 0 | 0 |
| NC_026438.1 Influenza A virus (A/California/07/2009(H1N1)) segment 1 polymerase PB2 (PB2) gene; complete cds | 0 | 0 |
| NJFV01000001.1 Pneumocystis jirovecii strain E2178 000000F; whole genome shotgun sequence | 0 | 0 |
| NZ_CP014267.1 Mycoplasma pneumoniae strain C267 chromosome; complete genome | 0 | 0 |
| NZ_LN831026.1 Corynebacterium diphtheriae strain NCTC11397 chromosome 1 | 0 | 0 |

0: no LAMP reaction

1: LAMP reaction occurs

Ref. 51.	Salinas, N. R.; Little, D. P., Electric LAMP: virtual loop-mediated isothermal AMPlification. *International Scholarly Research Notices*, 2012, (**2012**).
